# Data-Driven Strain Design Towards Mitigating Biomanufacturing Stresses

**DOI:** 10.1101/2023.09.17.558093

**Authors:** PV Phaneuf, SH Kim, K Rychel, C Rode, F Beulig, BO Palsson, L Yang

**Affiliations:** Novo Nordisk Foundation Center for Biosustainability, Technical University of Denmark, Kemitorvet, Building 220, 2800 Kongens, Lyngby, Denmark; Department of Bioengineering, University of California, San Diego, La Jolla, USA; Bioinformatics and Systems Biology Program, University of California, San Diego, La Jolla, USA; Department of Pediatrics, University of California, San Diego, La Jolla, CA, USA

**Keywords:** Genomics, ALE mutations, Data-driven strain design, Microbial cell factory, Reactive oxygen species, Acid stress, SOS response

## Abstract

Microbial strains used in large-scale biomanufacturing of melatonin often experience stresses like reactive oxygen species (ROS), SOS response, and acid stress, which can reduce productivity. This study leveraged a data-driven workflow to identify mutations that could improve robustness to these stresses for an industrially important melatonin production strain. This work integrated more than 7000 E. coli adaptive laboratory evolution (ALE) mutations to statistically associate mutated genes to 2 ROS tolerance ALE conditions from 72 available conditions. oxyR, fur, iscR, and ygfZ were significantly associated and hypothesized to contribute to fitness in ROS stress. Across these genes, 259 total mutations were inspected and 10 were chosen for reintroduction based on mutation clustering and transcriptional signals as evidence of fitness impact. Strains engineered with mutations in oxyR, fur, iscR, and ygfZ exhibited increased tolerance to H2O2 and acid stress, and reduced SOS response suggesting improved genetic stability. Additionally, new evidence was generated towards understand the function of ygfZ, a gene of relatively uncertain function. This meta-analysis approach utilized interoperable multi-omics datasets to identify targeted mutations conferring industrially-relevant phenotypes, describing an approach for data-driven strain engineering to optimize microbial cell factories.

**Visual Abstract:** 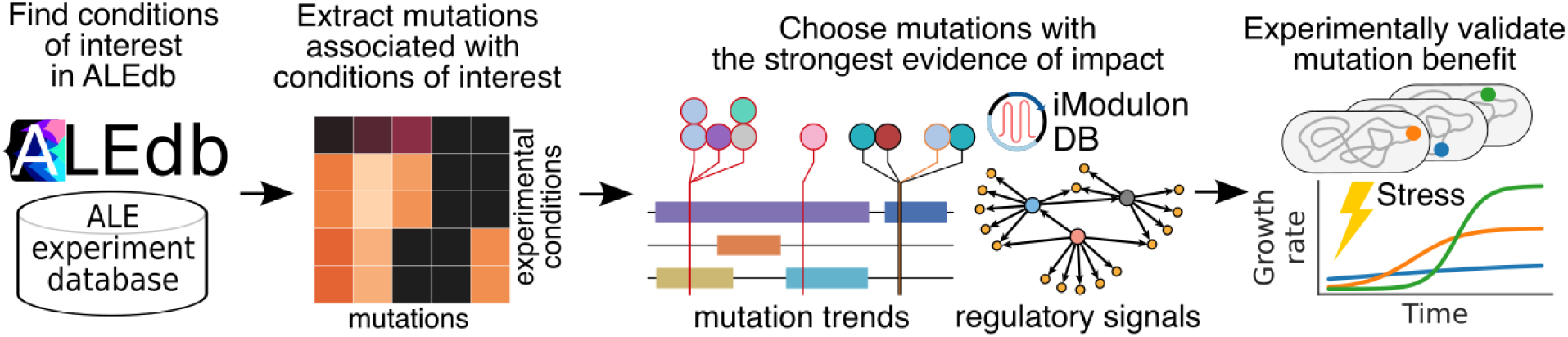

## Introduction

Strain robustness is critical for microbial cell factory development and bio-based manufacturing of chemicals or protein products. When microbial cells are cultivated in large volumes in bioreactors, they undergo many generations of cell divisions, and experience environmental stresses that can lead to poor strain performance. Here, we focus on stresses and responses that often occur in large-scale cultivation in bioreactors, including reactive oxygen species (ROS), acid stress, and the SOS response.

ROS, including H_2_O_2_, superoxide anion radical, and hydroxyl radicals, are toxic byproducts naturally generated during aerobic growth. ROS stress often occurs when oxygen level increases in the reactors, or when toxic substrates, intermediates or products accumulate, or NADPH depletes, etc. (1–3). When accumulated intracellularly, ROS can lead to damage of DNA, metalloproteins, and many cellular processes (3). Microorganisms develop various mechanisms to prevent ROS damage. The main transcriptional regulators involved in these processes include OxyR (oxidative stress regulator) that regulates redox balance and DNA/protein damage repair in response to hydrogen peroxide, Fur (Ferric Uptake Regulator) that is important for maintaining iron homeostasis, and IscR (Iron-Sulfur cluster Regulator) that regulates gene expression related to iron-sulfur cluster assembly and repair.

Severe ROS stress can lead to DNA damage through a variety of mechanisms including guanine oxidative lesions (4). This triggers SOS response, which is a series of responses to DNA damage. Strains that have elevated levels of SOS induction for an extended period of time may have decreased genetic stability and lose the desired production in fermentation (5). SOS response level can be an important indicator for strain robustness during fermentation scale-up.

Acid stress is another stress that might occur in fermentation. For example, in large-scale bioreactors, slow mixing of the base might lead to pH fluctuation. Acetic acid and amino acid accumulation could create a transient local low pH environment which is unfavorable in bioprocesses due to its negative impact on growth and production. Several studies have demonstrated the link between acid stress and ROS stress and further showed that engineered strains with reduced intracellular ROS can better survive at low pH (6–9). One of the reasons could be that iron sulfur clusters are labile to acid stress (10,11).

The rational design of strains for tolerance to adverse conditions is generally challenging due to unknown mechanisms involved (12). Adaptive Laboratory Evolution (ALE) is an experimental evolution method that has the potential to provide novel solutions to strain design in the form of mutations frequently not observed in natural isolates (13,14). ALEdb (aledb.org), a publicly accessible database of aggregated ALE mutations (15), has the potential to provide data that could lead to a more comprehensive understanding of gain-of-function mutations and result in a more effective strain design solution (16,17). Additional resources that are interoperable with ALEdb data are available for describing the consequences of ALE mutations, ultimately providing insights on which ALE mutations may be most effective for a desired phenotype. iModulons are independently-modulated gene sets computed from transcriptomic datasets, and their activities represent the abundances of co-regulated gene products under specific conditions (18). Comprehensive iModulon data is available through a publicly accessible database, iModulonDB ((imodulondb.org) (19,20). The combination of mutations and corresponding iModulon activity level changes in ALE strains have proven to be informative towards understand systems-level changes brought about by ALE mutations (21–25). The combination of ALE mutations and iModulon activities could also be used to focus mutation screening efforts on the subset of variants with the most promise of rendering phenotypes of interest. A recent study characterized mutations and iModulons from ALE strains that tolerated paraquat, a ROS-inducing agent, proposing some interesting hypothetical mechanisms which were not directly validated (26).

We have previously developed an *E. coli* strain producing melatonin (27). In this study, it was demonstrated that this strain has elevated ROS stress compared to the non-engineered ancestor strain. As a consequence, it also has a higher SOS response and decreased tolerance to acid stress. These changes can potentially result in poor performance in fermentation. In order to address this challenge, we developed a workflow that leverages ALE mutations to mitigate ROS stress, SOS induction, and acid stress. The resulting strains have decreased ROS and potentially improved strain robustness in fermentation scale-up.

## Results

### Production strain has elevated ROS stress and SOS response

We have previously engineered an *E. coli* strain producing melatonin by expressing heterologous enzymes required for melatonin synthesis from tryptophan (27), as well as genome engineering for improving tryptophan synthesis from glucose (Table 1, (28)). Since factors like product toxicity, acid accumulation and heterologous protein expression often lead to increased ROS stress (7,29), we tested the ROS stress level by growing the strain in H_2_O_2_. The melatonin production strain HMP3427 cannot grow in the presence of 10mM H_2_O_2_ in minimal medium after 72 h, suggesting that it is more sensitive to H_2_O_2_ compared to the wildtype.

**Table 1.** Strains and plasmids used in this study.

| ID | Genotype | Source |
| --- | --- | --- |
| pHM345 | P<sub>J23107</sub>-TpH(E2K, N97I, P99C)-PhhB-ASMT(A258E, V305A), Kan<sup>R</sup> | (46) |
| pSMART-SO<br>S-GFPuv | P<sub>cda</sub>-GFPuv, Kan<sup>R</sup> | (42) |
| pSD134 | P<sub>cda</sub>-GFPuv, Cm<sup>R</sup>, p15A | This work |
| pGE3 | *araC*, P<sub>ara</sub>-*gam-bet-exo*, tet<sup>R</sup>, P<sub>lacI</sub>-*tetO*, P<sub>tet</sub>-gRNA_pBR322, P<sub>J23105</sub>-MAD7, Amp<sup>R</sup>, SC101(ts) | (16) |
| pSD79 | P<sub>J23119</sub>-gRNA_Fur(P18T), Cm<sup>R</sup>, pBR322 | This work |
| pSD80 | P<sub>J23119</sub>-gRNA_YgfZ(T108P), Cm<sup>R</sup>, pBR322 | This work |
| pMA7 | P<sub>BAD</sub>-*bet-dam*, ColE1, Amp<sup>R</sup> | (39) |
| HMP3071 | BW25113 $\Delta tnaA \Delta trpR$ FolE(T198I) TrpE(S40F)<br>P<sub>FolE</sub>:P<sub>J23100</sub> $\Delta fhuA::P_2-ddc$ -P<sub>J23101</sub>-*aanat* | This work. Derived from previous work (27,46) |
| HMP3427 | HMP3071 + pHM345 | This work |
| DDB35 | BW25113 $\Delta fhuA$ | (47) |
| SDT392 | HMP3071 Fur(P18T) | This work |
| SDT764 | HMP3071 Fur(H71Y) | This work |
| SDT767 | HMP3071 Fur(R70S) | This work |
| SDT393 | HMP3071 YgfZ(T108P) | This work |
| SDT711 | HMP3071 IscR(V55L) | This work |
| SDT712 | HMP3071 OxyR(A213P) | This work |
| SDT713 | HMP3071 OxyR(A213T) | This work |
| SDT739 | HMP3071 Fur(H71Y) YgfZ(L29R) | This work |
| SDT744 | HMP3071 Fur(H71Y) YgfZ(L29R) OxyR(A213T) | This work |

We further tested the SOS response level in the production strain since ROS stress can lead to elevated DNA damage and SOS response. We used an SOS biosensor that was previously reported (30). As shown in Figure 1, HMP3427 showed a slightly higher SOS response in normal growth conditions. When treated with H_2_O_2_, the difference in SOS response levels between HMP3427 and wild type increased dramatically. These data suggest that mitigating ROS stress could potentially improve strain robustness and performance.

**Figure 1.**
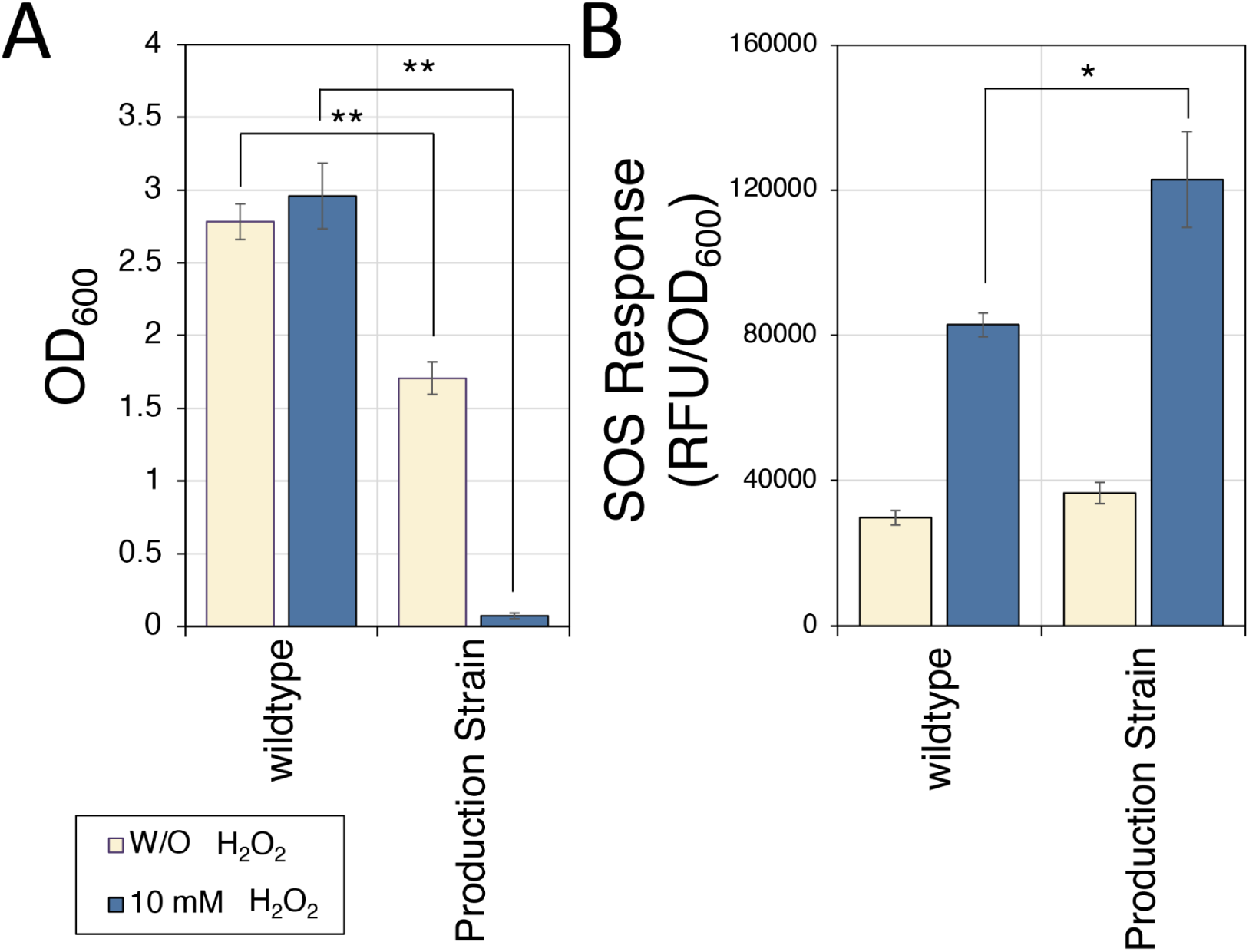
Melatonin production strain has higher ROS stress sensitivity and SOS response compared to the wildtype. (A) Growth of wildtype strain and melatonin production strain with or without H_2_O_2_. The wildtype strain was able to tolerate 10mM H_2_O_2_ but the production strain cannot. (B) SOS response of the wildtype and melatonin production strain, measured by a GFP-based SOS sensor. The production strain has a slightly higher SOS response when growing in LB medium, and about 50% higher SOS response level in the presence of H_2_O_2_ (*p < 0.05, **p < 0.005). See materials and methods for details.

### A data-driven approach to identify mutations that benefit ROS stress tolerance

To identify variants that could render ROS tolerance to a host, a meta-analysis was performed on adaptive laboratory experiment mutations, their conditions, and their potential impact. Evidence of the impact for individual mutations on resulting mutated features of interest, such a frequency of mutation and substantial transcriptional changes, was then used to determine which mutation was most likely to provide maximum effect and potential benefit. Finally, a small subset of mutations for each genetic feature of interest was chosen for reintroduction and testing of increased ROS tolerance.

The primary datatypes and evidence impact for individual mutations were ALE mutations, their conditions, and the iModulon activities for the samples hosting ALE mutations of interest. ALE mutations are variants that manifest during ALE experiments and were exported from ALEdb (aledb.org) (15). iModulons are independently-modulated gene sets and their activity levels represent the abundances of their gene products relative to specific conditions (20). iModulon data was exported from imodulondb.org (19,20).

ALEdb public data was used to find significant associations between mutated genetic features and ROS agents (7680 public mutations across 72 unique conditions) (Figure 2A). mutations *oxyR* was the single genetic feature statistically associated with both sources of ROS stress (paraquat, FeSO_4_), ignoring other shared mutations which are frequently observed in ALE and likely have more general benefits (*rph*, *pyrE*/*rph*) (Figure 2A). Of the remaining statistically associated mutated features, mutations to *fur*, *iscR*, and *ygfZ* have been proposed as beneficial for general ROS-stress tolerance (26).

**Figure 2.**
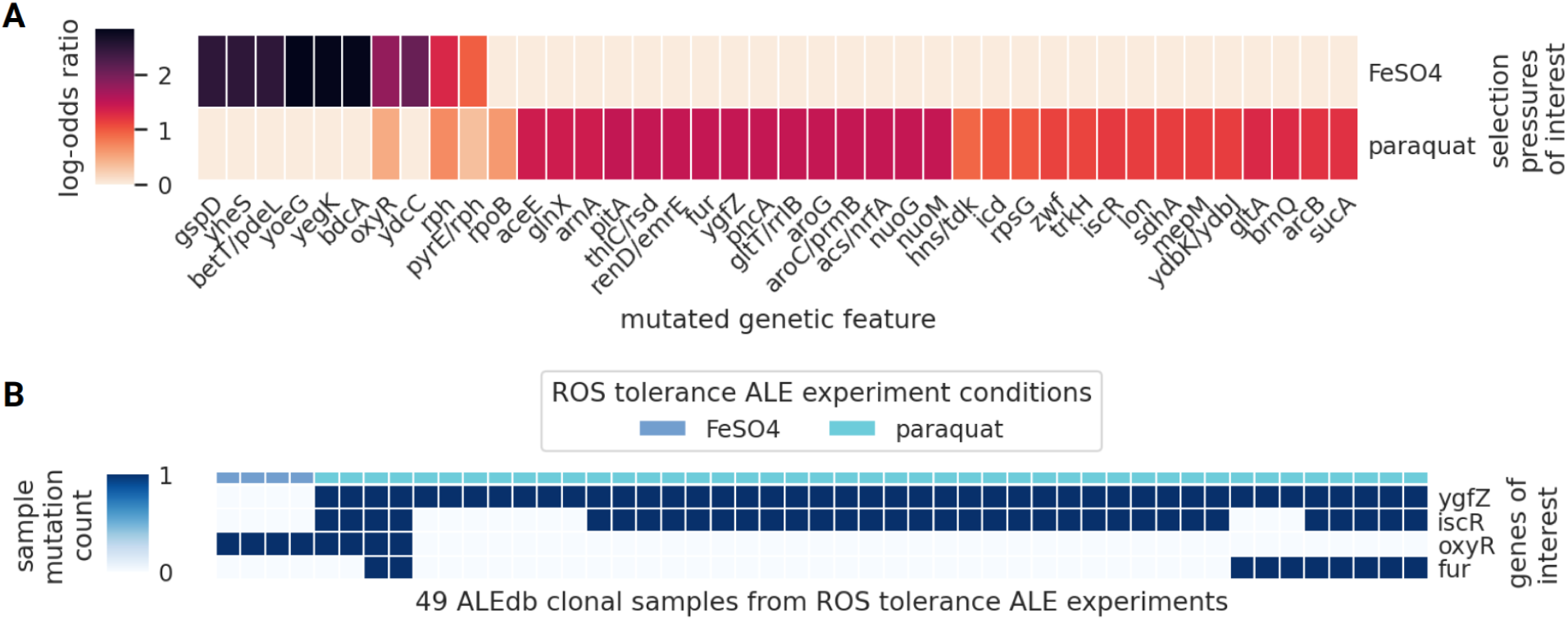
Meta-analysis of mutated genetic features and their experimental conditions in ALEdb samples. A) ALEdb public mutated genetic features statistically associated with sources of ROS stress (Fisher’s Exact Test, p-value < 0.01, Bonferroni corrected). B) The mutation count of genes of interest in samples for ALE experiments explicitly involving ROS stresses.

The co-occurrence of mutated genes of interest within the same strain was investigated to understand if a combination of mutations was feasible, if not more beneficial. Most samples exposed to paraquat had more than one gene of interest mutated, with some samples having all 4 (Figure 2B).

*oxyR* mutants were associated with both ALEdb ROS stress conditions of paraquat and FeSO4 (Figure 2) and 102 public and private mutations to *oxyR* were extracted from ALEdb for *oxyR* (Figure 3A). Mutations to *oxyR* were proposed to activate ROS protective and preventative functions regulated by OxyR and strains evolved in conditions involving ROS stressors and hosting *oxyR* mutations had higher growth rates (25,26). ALE mutations were found in both of OxyR’s 2 major domains: the DNA and substrate binding domain (Figure 3). The majority of mutations to OxyR result in nonsynonymous substitutions, though there exists mutations resulting in amino acid deletions or premature truncations (Figure 3A). The substrate binding domain, hosting the OxyR subunit interface and disulfide bond sites that mutations generally cluster around, host all of the ROS-related mutations (Figure 3), emphasizing this area as being important for ROS-related selection pressures. An amino acid position in this area, A213, was mutated in both ALEdb ROS-related selection pressures (Figure 3A).

**Figure 3.**
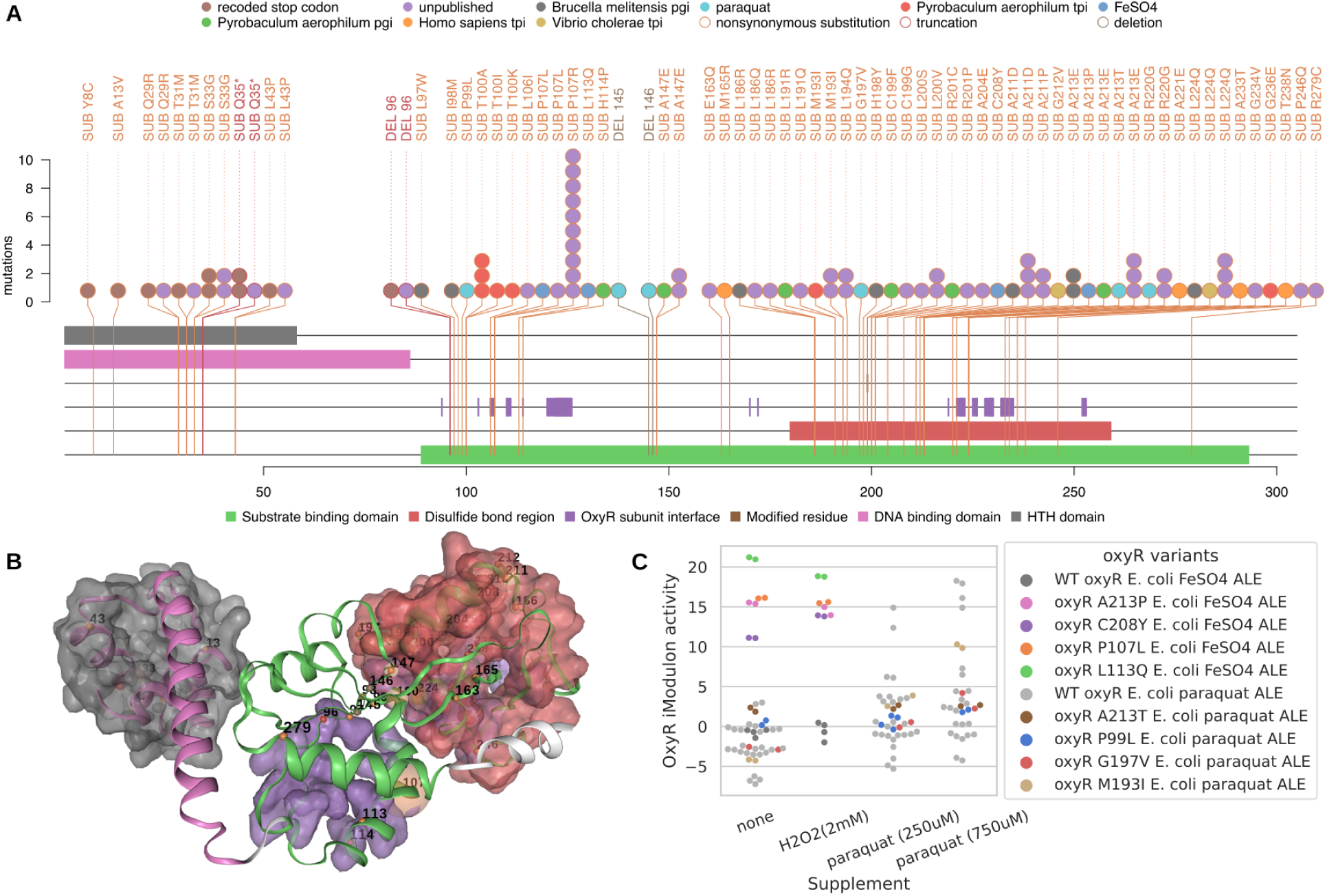
ALEdb mutations, their effects on OxyR, and to iModulons. A) Mutation needle plot demonstrating the effect and position of ALEdb mutations to *oxyR*. B) OxyR’s 3D structure and mutated residues from mutations. The residue chain and transparent surfaces are colored according to the legend of the corresponding mutation needle plot. Mutations are represented by a small opaque sphere with a value representing their amino acid position on the corresponding mutation needle plot. The color of the mutation’s sphere corresponds to the mutation’s predicted effect as described by the legend on the corresponding mutation needle plot. The transparent sphere centered on the mutations’ opaque sphere represents the number of mutations with a specific predicted effect on that position. C) OxyR iModulon activity for strains involved in ALE experiments with paraquat or high FeSO_4_ as ROS selection pressures.

OxyR is a transcription factor regulating genes of the OxyR iModulon, which respond to oxidative stress (31). iModulon data was available for 4 *oxyR* mutations from the ALE experiments involving FeSO_4_ and 3 *oxyR* mutations from the ALE experiment involving paraquat (19,20). The mutation strains were endpoints of ALE experiments that had a mutation in *oxyR* selected by ALE as well as others specific to each evolution. Along with other endpoint strains from these ALE experiments, the *oxyR* mutation strains were subjected to either FeSO_4_ or paraquat, depending on their original ALE experiment, and their OxyR iModulon activities were determined (Figure 3C). FeSO_4_ ALE *oxyR* mutations activated the OxyR iModulon across all conditions they were subjected to; this is possibly due to epistasis with other ALE endpoint mutations. All mutants from the paraquat ALE except for *oxyR* A213P generally increased their OxyR iModulon activity with an increase in paraquat, indicating that A213P was the only mutant with consistent OxyR iModulon activation. Due to both ROS ALE experiments selecting *oxyR* A213 mutants and A213 mutations demonstrating consistent substantial OxyR iModulon activity in all conditions, both A213T and A231P mutations were proposed for reintroduction (Table S1).

*fur* mutants were associated with the ROS stress condition of paraquat (Figure 2) and 39 public and private mutations to *fur* were extracted from ALEdb (Figure 4A). ALE mutations were found in both of Fur’s 2 major domains (Figure 4A, Figure 4B). These mutations demonstrated 3 trends of interest: 1) all of the paraquat mutations are hosted on the DNA binding region, 2) mutations cluster in the first half of Fur’s amino acid sequence, and 3) mutations manifest on or near the subunit interface (Figure 4A). On Fur’s 3D structure, it becomes more clear that mutations to residues 7, 14 18, 23, and 42 comprise a cluster that doesn’t seem to target the subunit interface; besides mutations to residues 102 and 104, the remaining mutations were found in close proximity to subunit interfaces (Figure 4A, Figure 4B).

**Figure 4.**
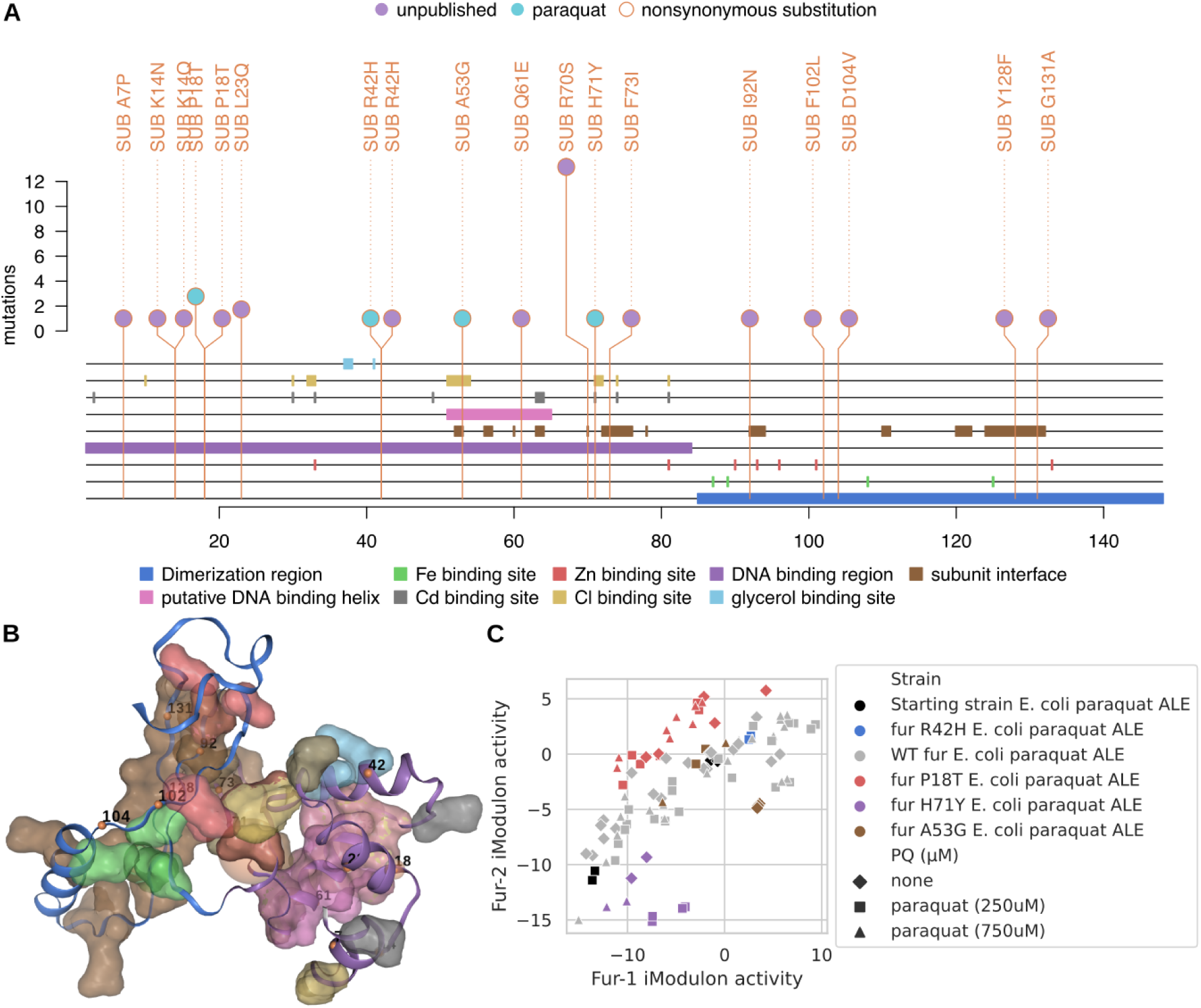
ALEdb mutations and their effects to Fur. A) Mutation needle plot demonstrating the effect and position of ALEdb mutations to *fur*. B) Fur’s 3D structure and mutated residues from mutations. The residue chain and transparent surfaces are colored according to the legend of the corresponding mutation needle plot. Mutations are represented by a small opaque sphere with a value representing their amino acid position on the corresponding mutation needle plot. The color of the mutation’s sphere corresponds to the mutation’s predicted effect as described by the legend on the corresponding mutation needle plot. The transparent sphere centered on the mutations’ opaque sphere represents the number of mutations with a specific predicted effect on that position. C) Fur-1 and Fur-2 iModulon activity for samples with *fur* mutations from an ALE experiment with paraquat as the selection pressure.

Fur regulates 2 iModulons that are associated with ferric uptake: Fur-1 and Fur-2 (26). Fur-1 primarily describes systems of siderophore synthesis and transport while Fur-2 describes iron and siderophore transport systems as well as hydrolysis systems (26). iModulon data was available for paraquat ALE endpoints containing *fur* mutations P18T, H71Y, R42H and A53G exposed to different concentrations of paraquat (Figure 4C) (19,20). Fur-1 and Fur-2 iModulons activities vary across samples, though demonstrated a trend (26) (Figure 4C), where *fur* P18T mutations were consistently above the trend, corresponding with a general increase in Fur-2 activity, and H71Y mutations were consistently below the trend, corresponding with a general decrease in Fur-2 activity. P18T is found in the cluster of mutations not targeting subunit interfaces and therefore could represent this set and their potential of decreasing Fur-2 iModulon activity. H71Y is part of the set of mutations landing near or on subunit interfaces and could represent this set and their potential for increasing Fur-2 iModulon activity. R70S manifests an order of magnitude more than H71Y in ALEdb and may have similar effects. All three of these mutations were proposed for reintroduction to represent all observed trends (Table S1).

*iscR* mutants were associated with the ROS stress condition of paraquat (Figure 2) and 72 public and private mutations to *iscR* were extracted from ALEdb (Figure 5). ALE mutations generally clustered on or near 2Fe-2S binding sites, subunit interfaces, the DNA binding region, and the HTH domain (Figure 5). Mutations generally clustered in 2 areas on both the AA sequence (Figure 5A) and protein structure (Figure 5B); mutations in each cluster land on or near unique functional annotations as well subunit interface sites.

**Figure 5.**
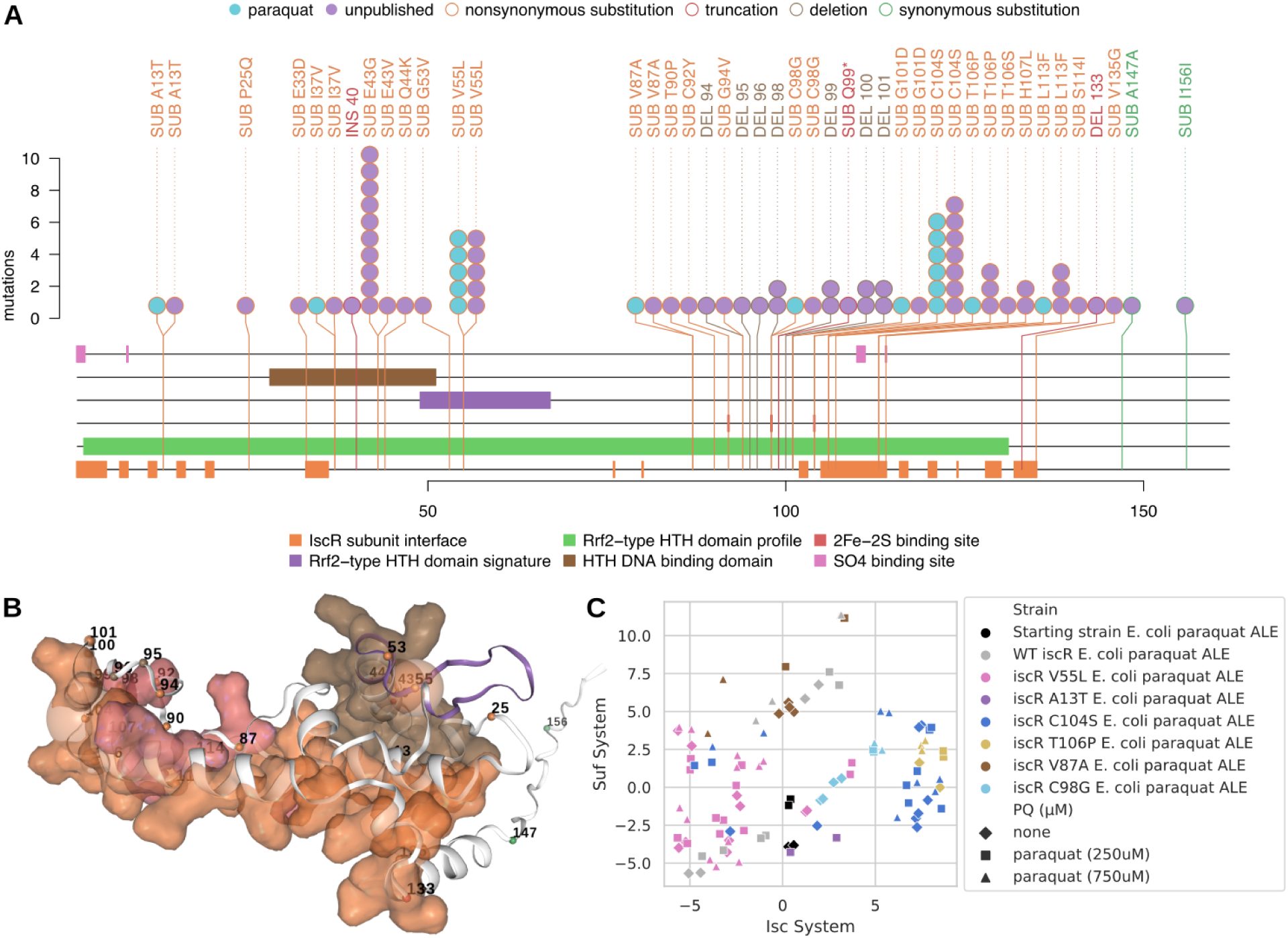
ALEdb mutations and their effects to IscR. A) Mutation needle plot demonstrating the effect and position of ALEdb mutations to *iscR*. B) IscR’s 3D structure and mutated residues from mutations. The residue chain and transparent surfaces are colored according to the legend of the corresponding mutation needle plot. Mutations are represented by a small opaque sphere with a value representing their amino acid position on the corresponding mutation needle plot. The color of the mutation’s sphere corresponds to the mutation’s predicted effect as described by the legend on the corresponding mutation needle plot. The transparent sphere centered on the mutations’ opaque sphere represents the number of mutations with a specific predicted effect on that position. C) Isc and Suf iModulon activity for samples with *iscR* mutations from an ALE experiment with paraquat as the selection pressure.

IscR regulates the Isc and Suf iModulons that are both associated with iron-sulfur cluster synthesis (26). iModulon data was available for paraquat *iscR* ALE mutations A13T, V55L, V87A, C98G, C104S, and T106P (Figure 5C) (19,20). Each mutation seemed to correspond with trends in increasing or decreasing activities of either or both Suf and Isc iModulons. C104S and C98G corresponded with increasing Isc activity. V87A corresponded with increasing Suf activity. A13T corresponded primarily with decreasing Suf activity. Finally, V55L and T106P corresponded with decreased Isc activity, and in the case of V55L, also decreased Suf activity. V55L is also the only mutation with iModulon data found in the cluster near the DNA binding region and HTH domain; all others are found in the cluster near the 2Fe-2S binding sites, SO4 binding sites, and/or subunit interfaces. For mutations that corresponded with higher Isc iModulon activity, C104S consistently corresponded with the highest activity. V55L consistently corresponded with the lowest Isc and Suf activity. V87A generally corresponded with the highest Suf activity. These mutations were therefore considered representative for their effects on iModulons and were proposed for reintroduction (Table S1).

YgfZ is a folate-binding protein that plays a role in iron-sulfur assembly or repair (32). Its function has not been well characterized yet, but there is evidence that it is important for oxidative stress resistance (Figure 2A). *ygfZ* mutants were associated with the ROS stress condition of paraquat (Figure 2A) and 46 public and private mutations to *ygfZ* were extracted from ALEdb (Figure 6). ALE mutations generally clustered on or near 2 functional annotations: 1) the Aminomethyltransferase folate-binding domain, and 2) the GcvT family signature motif (Figure 6A). On YgfZ’s 3D structure, mutation clusters are more suggestive of 2 planes, with one plane targeting ethandiol binding sites and the other sitting between all binding sites of the Aminomethyltransferase folate-binding domain and the GcvT family signature motif (Figure 6B). Of all these mutations, L29R, V107E, and T108P were most frequently mutated in the paraquat ALE experiment and belonged to 2 different 1D clusters around the Ethandiol binding sites.

**Figure 6.**
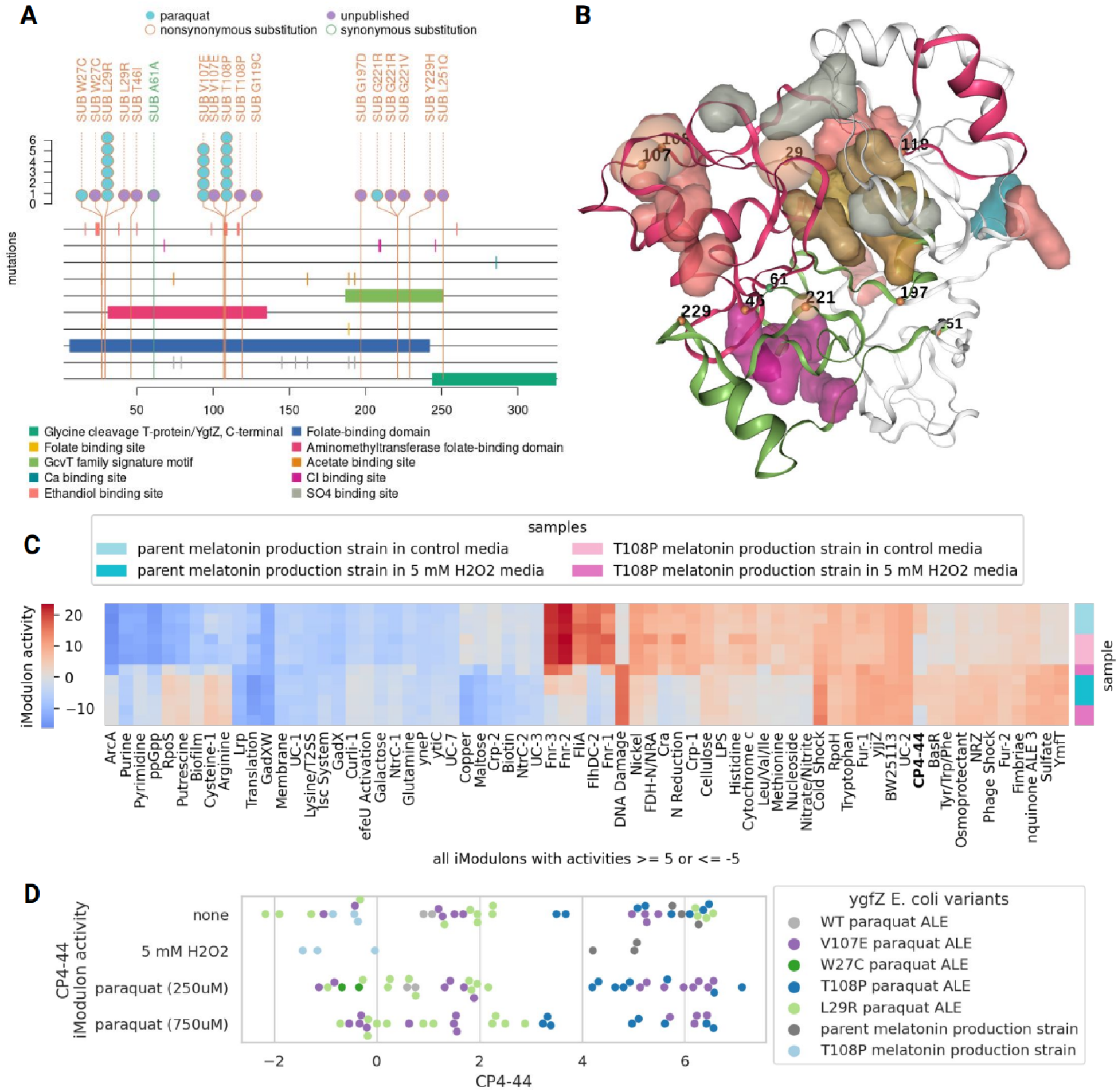
Mutation needle plot demonstrating the effect and position of ALEdb mutations to YgfZ. A) Mutation needle plot demonstrating the effect and position of ALEdb mutations to fur. B) YgfZ’s 3D structure and mutated residues from mutations. The residue chain and transparent surfaces are colored according to the legend of the corresponding mutation needle plot. Mutations are represented by a small opaque sphere with a value representing their amino acid position on the corresponding mutation needle plot. The color of the mutation’s sphere corresponds to the mutation’s predicted effect as described by the legend on the corresponding mutation needle plot. The transparent sphere centered on the mutations’ opaque sphere represents the number of mutations with a specific predicted effect on that position. C) A heatmap of iModulon activities for samples from a *ygfZ* mutant characterization experiment using the melatonin production strain as the base strain. D) The CP4-44 iModulon activity for multiple strains across conditions involving oxidative stress.

In order to better understand the function of YgfZ, iModulon activities derived from transcriptional profiles of the melatonin production strain with or without the T108P YgfZ variant were compared in both normal conditions and with H_2_O_2_ treatment (Materials and Methods). Surprisingly, the T108P YgfZ variant had little impact on the iModulon activity relative to the presence of H_2_O_2_, though did coincide with different CP4-44 iModulon activity regardless of environmental conditions (Figure 6C). The CP4-44 iModulon, which contains most of the genes for the CP4-44 prophage, was consistently deactivated in the presence of the T108P YgfZ variant. After this observation, all iModulonDB *E. coli* samples with *ygfZ* variants were gathered and their CP4-44 iModulon activities compared. In *E. coli* ALE endpoint strains exposed to paraquat, the L29R and W72C YgfZ variant consistently coincided with reduced CP4-44 iModulon activity. The T108P YgfZ *E. coli* melatonin production mutant consistently coincided with reduced CP4-44 iModulon activity in both H_2_O_2_ treatment and no treatment relative to the parent *E. coli* melatonin production strain. L29R and T108P were therefore chosen for reintroduction due to their high frequency of manifestation in ALE experiments, location in different variant clusters, and their evidence of reduce CP4-44 iModulon activities (Table S1).

### ALEdb mutations increased tolerance to ROS-related stresses

We constructed 9 strains reintroducing single or combinations of ALE mutations (Table S1, Figure 7) using CRISPR-MAD7 or MAGE (Materials and Methods). Combinations were pursued due to the potential for synergistic effects between mutations as evidenced by their co-occurrence in ALEdb clonal samples (Figure 2B). Among all the mutations, YgfZ(L29R) and OxyR(A213T) only appeared in the multiplex MAGE isolates in combination with Fur(H71Y). We failed to construct IscR(V87A) and IscR(C104S) using both MAGE and CRISPR-MAD7, suggesting that these mutations might have growth defect in our strain and the benefit of these mutations rely on the presence of other ALE mutations. We tested the growth of 9 strains in glucose minimal medium with or without H_2_O_2_ (Figure 7A). While the production strain hardly grew under 10 mM H_2_O_2_, in contrast, a subset of mutants demonstrated more substantial growth. Strains containing Fur(P18T), Fur(H71Y), Fur(R70S) and OxyR(A213T) mutations had inconsistent growth among the four biological replicates. The strain containing OxyR(A213P) reached the same OD consistently with or without 10mM H_2_O_2_, suggesting that this mutation offers superior benefit towards ROS resistance.

**Figure 7.**
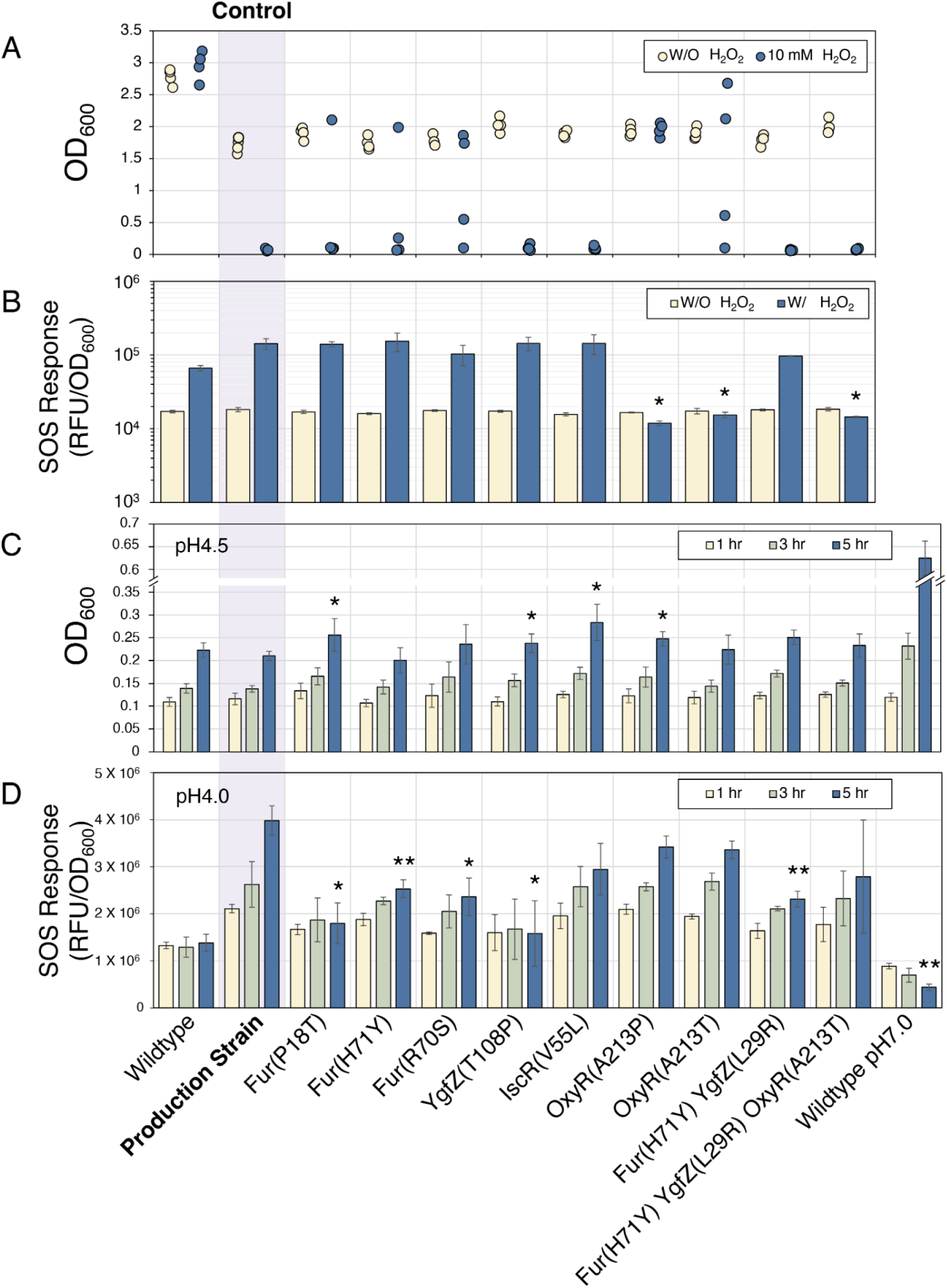
Comparison of growth and stress response between the ancestor melatonin production strain (control) and the production strains with ALE variants in H_2_O_2_ and acid stress. A) Biomass represented by OD (600 nm) of wildtype, melatonin production strain, and strains with ALE mutations implemented after H_2_O_2_ treatment for 72 h (Materials and Methods). B) SOS response of wildtype, melatonin production strain and production strains with ALE variants with or without 10 mM H_2_O_2_ treatment. Data represents the average of 3 replicates. Error bars indicate standard deviation (*p < 0.05). C) Tolerance of melatonin production strain and ALE variants in acid stress. Cultures of neutral pH were diluted into pH 4.5. OD (600 nm) was monitored after 1 h, 3 h, and 5 h (Materials and Methods). The height of the bars indicate the average concentration of 3 biological replicates and error bars indicate the standard deviations (*p < 0.05). D) SOS response of melatonin production strain and ALE variants in acid stress. Cultures of neutral pH were diluted into pH 4.0. SOS response was monitored after 1 h, 3 h and 5 h using a GFP sensor (Materials and Methods). The height of the bars indicate the average of the 3 biological replicates. The error bars indicate the standard deviations. The dots represent the OD values of each of the replicates (*p < 0.05, **p < 0.005).

We also examined the SOS response of the ALE mutations. All the strains were transformed with the SOS reporter plasmid pSD134. The resulting strains were cultivated in glucose minimal media with or without H_2_O_2_ for 24 h. After the treatment of H_2_O_2_, the production strain showed a very high SOS response compared to wildtype strain (Figure 7B). Among all the ALE mutations, the strains containing Fur(R70S) and Fur(H71Y) + YgfZ(L29R) demonstrated reduced SOS response compared to the original production strain. Surprisingly, the 3 strains containing OxyR(A213P) and OxyR(A213T), either in singleton or in combination with other mutations, did not show elevated SOS response under the stress of ROS species at all (Figure 7B). This suggested that OxyR(A213P) and OxyR(A213T) mutations have activated ROS tolerance machinery and prevented SOS response caused by H_2_O_2_.

We further tested if the ALE mutations granted any benefits in acid stress, as ROS stress mitigation can be beneficial for acid tolerance. We grew the cells in glucose M9 minimal media at pH 7.0 overnight and diluted the cultures into pH 4.5 to OD_600_ 0.1. We monitored the OD after 1 h, 3 h and 5 h. As shown in Figure 7C, the wildtype strain is still able to maintain the biomass after 5 h exposure in pH 4.5, however the production strain has a slight decrease of OD. Variants containing Fur(P18T), YgfZ(T108P), IscR(V55L), OxyR(A213T) have significantly improved survival in low pH.

We also tested the effect of ALE mutations in SOS response during acid stress (Figure 7D). We performed similar cultivation experiments using strains containing the SOS sensor plasmid. We grew the strains in normal M9 glucose medium (pH 7.0) overnight. Then we transferred the cultures into the new M9 glucose media pH 4.0 to OD_600_ 0.05. We took samples after 1 h, 3 h and 5 h and measured fluorescence. As shown in Figure 7D, all ALE mutations are beneficial in reducing SOS response during acid stress. Noticeably, strains containing Fur(P18T) or YgfZ(T108P) have lower SOS response than other variants, indicating that these two mutations have a stronger benefit managing the acid stress.

## Discussion

This study described a data-driven workflow leveraging both *E. coli* ALE mutations and iModulon activities to identify a small set of mutations that had evidence of potentially conferring ROS tolerance. Strains incorporating a subset of these mutations were found to have tolerance not only to ROS stress but also to acid stress and reduced SOS responses. These results have several primary implications.

First, interoperable ALE mutation and iModulon data types provided valuable evidence for successfully identifying mutations with substantial physiological impact in the presence of ROS stress. This evidence was enriched through the aggregation of data from a variety of experiments and revealed mutations from different experiments that may have a similar impact. Functional annotations for genes were aggregated from multiple sources and hinted at what gene product functions were targeted by mutations. iModulon data proved valuable in interpreting the potential magnitude of impact a mutation may have as well as initial suggestions of the systemic effects of the mutations. The iModulon data enabled screening efforts to focus only on the mutations with the strongest evidence of substantial physiological impact, rather than having to rescreen the much larger set of all available mutations. The results of this study demonstrate the value of combining aggregated and interoperable data types towards strain design and this strategy could be reused for similar efforts.

Second, most of the selected ALE mutations offer benefits towards H_2_O_2_ or acid stress. All OxyR variants we have tested have significantly increased H_2_O_2_ tolerance, which is expected as OxyR is known to be more sensitive towards H_2_O_2_ (33). Both A213T and A213P mutations reduced SOS response in H_2_O_2_, while A213P is the only mutation that offered consistent growth in H_2_O_2_.

Acid stress is another stress that is relevant to industrial processes. Cells respond to acid stress using different mechanisms, including decarboxylation reactions and ion transporters to reduce protons (34). Internal pH drop can cause damage to DNA and iron-sulfur clusters (10,35). In a separate transcriptomic analysis, many genes activated upon acid stress largely fall into OxyR and SoxR regulations (36). Fur(P18T), YgfZ(T108P), IscR(V55L), OxyR(A213T) mutations all showed improved tolerance towards acid stress, indicated by higher OD and lower SOS response (Figure 7C, D). Fur(P18T) and YgfZ(T108P) mutations both had the lowest SOS response while manifesting some of the highest growth in acid stress.

YgfZ(T108P) showed surprisingly low impact on the transcriptome of *E. coli* in normal growth (M9 glucose medium) or with H_2_O_2_ stress. The iModulon representing prophage CP4-44 deactivation is the only iModulon with substantially different activities coinciding with the presence or absence of YgfZ(T108P). YgfZ(T108P) was shown to coincide with a significant reduction of the SOS response in acid stress (Figure 7D). Prophage gene expression is thought to be triggered by the SOS response (37). We speculate that YgfZ(T108P) reduced the SOS response in some form, which subsequently reduced the activity of the CP4-44 iModulon. Exactly how YgfZ(T108P) lowers CP4-44 prophage activity requires further investigation.

Third, commonly used approaches to mitigate ROS stress include increasing redox cofactors NADPH. For example, genetic manipulations that boost oxidative pentose phosphate (PP) pathway and/or reduce upper glycolic flux are efficient ways to improve NADPH pool upon oxidative stress. However these strategies can be complicated to implement and sometimes lead to growth defects (3,38). The mutations we present in this work uses different mechanisms, providing an easy alternative to address the ROS stress. For example, YgfZ(T108P) mutation resulted in very little perturbation to the transcriptome compared to the parent strain (Figure 6), which can be advantageous in many applications.

We also tested strains containing Fur(P18T), YgfZ(T108P), and OxyR(R213P) in batch fermentation (Supplementary Information). We did not observe significant improvement in the mutant strains, all the three mutant strains produced similar titers to the parent strain (Figure S1). It will be valuable to test the mutant strains in conditions where ROS stress tolerance and strain robustness are more relevant, such as large scale bioreactors with stresses caused by spatial heterogeneity and cultivation processes where genetic instability can lead to poor-performing variants.

The workflow and findings from this study have important broader implications for the field of strain engineering. First, as increasingly large datasets become available, meta-analysis approaches leveraging multiple data types will become more valuable for identifying target genes or mutations. Second, the ability to pinpoint a small set of mutations that improve robustness without compromising production highlights the potential of data-driven design. Compared to traditional approaches that perturb central metabolism, mutations identified through multi-omics evidence may enable more targeted improvements. Finally, the redesigned strains exhibiting higher stress tolerance underscores the promise of integrating systems-level data for engineering industrially-relevant phenotypes. With further testing, strains designed using this broadly-applicable workflow could provide enhanced productivity and stability for large-scale biomanufacturing processes. Overall, this study provides a framework for data-driven strain engineering that could accelerate the design of optimized industrial hosts.

## Materials and Methods

### Strain construction

The *E. coli* strain HMP3427 is derived from a previously reported melatonin production strain which is based on BW25117 (Table 1). All the ALE mutations were implemented into the genome of HMP3271 (background strain of HMP3427) using one of the following methods. SDT392 and SDT393 were generated by CRISPR/MAD7 as described previously (16). SDT711, SDT712, and SDT713 was generated through single target TM-MAGE (39) where repair oligos contain only one editing target. Other variants were created using TM-MAGE where repair templates contain a library of oligos. We performed 2 rounds of transformation of a pool of MAGE oligos (4 μL, pre-mixed in a tube with a total concentration of 100 nmol/ml). Each 48 isolate was analyzed by illumina genome sequencing to validate mutations on the genome. Clones containing single (SDT764, SDT767) or multiple mutations (SDT739, SDT744) were selected for further analysis. All plasmids used in this study were constructed using USER cloning (40).

### Growth test under ROS stress

Melatonin production plasmid (pHM345) was transformed into each ALE variant by chemical transformation and spreaded on Luria-Bertani (LB) agar plate containing 50 μg/L of kanamycin. Followed by overnight incubation at 37°C, four colonies of each strain were inoculated in 300 μL of Luria-Bertani (LB) liquid medium supplemented with kanamycin (50 μg/L) in 96-deepwell plate and incubate at 37°C for overnight with shaking at 250 rpm. 10 μL of each cultivation was transferred into 250 μL of M9 minimal medium containing 2 g/L of glucose with or without 10 mM H_2_O_2_, and subjected to Growth Profiler (Enzyscreen, Heemstede, Netherlands) to monitor the growth at 30°C with 250 rpm for 72 h. Growth rates were calculated by the Croissance package (41). Kanamycin was not added for DDB35, control strain in the medium.

### SOS response sensor assay

SOS response sensor protein was obtained from addgene (pSMART-SOS-GFPuv, Plasmid #102283) (42) and cloned into a backbone containing p15A origin and chloramphenicol resistance to construct pSD134. Each strain was co-transformed with melatonin production plasmid (pHM345) and pSD134. Strains were cultivated in triplicate into 300 μL of Luria-Bertani (LB) liquid medium supplemented with chloramphenicol (25 μg/L) and kanamycin (50 μg/L) in 96-deepwell plate at 37°C for overnight. Each 10 uL of broth was transferred into 300 μL of LB medium supplemented with chloramphenicol (25 μg/L) and kanamycin (50 μg/L) in a 96-deepwell plate. Plate was duplicated and incubated in a 37°C shaking incubator at 250 rpm to reach OD_600_ 0.5. 10 mM of H_2_O_2_ was added into one plate and the fluorescence was monitored after 2h (Figure 1) or 24 h (Figure 7B) (excitation; 485 nm, emission; 520 nm).

### Acid Stress Assay

For SOS response monitoring, each strain was co-transformed with pHM345 and pSD134. Three independent colonies of each strain were cultivated in 300 μL of M9 minimal medium (pH7.0) containing 0.2% glucose, chloramphenicol (25 μg/L) and with kanamycin (50 μg/L) for pHM345 containing strains in 96-deepwell plate at 37°C overnight. Cells grown overnight were transferred into 400 μL of the same medium adjusted to pH 4.0 to start the cultivation at OD_600_ 0.05 in 96-deepwell plate and incubated in a 37°C shaker. OD_600_ and fluorescence were monitored after 1 h, 3 h, and 5 h as described above. For growth monitoring of each strain, pHM345 was transformed into background strains. Six independent colonies of each strain were picked and inoculated in 300 uL of M9 minimal medium (pH7.0) containing 0.2% glucose and kanamycin (50 μg/L) in 96-deepwell plate at 37°C overnight. Cells grown overnight were transferred into 400 μL of the same medium adjusted to pH 4.5 to start the cultivation at OD_600_ 0.1 in 96-deepwell plate and incubated in a 37°C shaker. OD_600_ was monitored after 1 h, 3 h, and 5 h.

### DNA Resequencing

All strains implemented with ALE mutations were sequenced and validated in house (CfB Biofoundry). For genomic DNA of *E. coli* samples, CyBio Felix robot and the smart DNA prep (a96) – FX kit (Analytik Jena) was used to extract genomic DNA. PlexWell 384 kit (SeqWell) was used for tagmentation, barcoding and library amplification. The final library pool was sequenced with NextSeq. Variant data was generated through ALEdb, which uses the breseq mutation finding pipeline (43,44). Being that these samples come from different projects, various versions of breseq were used in their mutation data generation. The sequencing reads used to generate the mutation data were subjected to quality control through either FastQC (https://www.bioinformatics.babraham.ac.uk/projects/fastqc/) and the FastX-toolkit (http://hannonlab.cshl.edu/fastx_toolkit/) or AfterQC (45).

### RNA sample preparation

HMP3427 and SDT495 were streaked on the LB agar plate containing 50 μg/L of kanamycin and incubated at 37°C overnight. Seed culture was made by inoculating each three colonies into 5 mL (in 50 mL tube) of M9 minimal medium supplemented with 4 g/L of glucose and 50 μg/L of kanamycin followed by shaking at 250 rpm and 37°C. OD_600_ of each seed culture was measured after overnight incubation and adjusted to 0.05 into 25 mL of the same medium (in 250 mL baffled flask) to start main culture, followed by incubation at 30℃ and 250 rpm. Each main culture was duplicated to compare the effect of H_2_O_2_ treatment. Growth was monitored by measuring OD_600_ until they reached 0.5. Then, 5 mM of H_2_O_2_ was added to three flasks of each strain, and they were further incubated for 2 h. Culture broths were taken and transferred into each tube containing two volumes of RNAprotect Bacteria Reagent (Qiagen, Hilden, Germany). Samples were pelleted as manufacturer’s instructions and stored at −80°C freezer until extraction. RNAs were extracted using QIAcube (Qiagen, Hilden, Germany) with the RNeasyProtectBacteria/BacterialCellPellts/DNaseDigest protocol. Sequencing of RNA samples was performed by Azenta Life Sciences (Burlington, Massachusetts, US) with NEBNext®Ultra™ II Directional RNA Library Prep Kit for Illumina (NEB, Ipswich, Massachusetts, US) for library preparation.

### Generative AI

ChatGPT, BARD, Claude.ai, and Grammarly were used to refine the narrative and grammar of this work, though were not used in the research process.

## Supplementary information

**Supplementary Table S1.** Table of mutations reintroduced and tested for increased fitness in the presence of ROS agents.

| Gene | Variant | Reasons for Inclusion | Source |
| --- | --- | --- | --- |
| *fur* | P18T | Hypothesized to increase iron uptake to counteract higher ROS levels | (26) |
| *fur* | H71Y | Hypothesized to decrease iron uptake to prevent iron toxicity | (26) |
| *fur* | R70S | Unpublished ALEdb variant near H71Y | Unpublished ALEdb variant |
| *iscR* | C104S | Hypothesized to rebalance Fe-S cluster synthesis and SoxS for better ROS readiness | (26) |
| *iscR* | V87A | Hypothesized to rebalance Fe-S cluster synthesis and SoxS for better ROS readiness | (26) |
| *iscR* | V55L | Hypothesized to rebalance Fe-S cluster synthesis and SoxS for better ROS readiness | (26) |
| *ygfZ* | T108P | Putative Fe-S cluster repair gene; hypothesized to reduce SOS response | (26) |
| *ygfZ* | L29R | Putative Fe-S cluster repair gene; hypothesized to reduce SOS response | (26) |
| *oxyR* | A213T | A213 mutated in multiple ROS studies | (25, 26) |
| *oxyR* | A213P | A213 mutated in multiple ROS studies | (25, 26) |

**Supplementary Figure S1.**
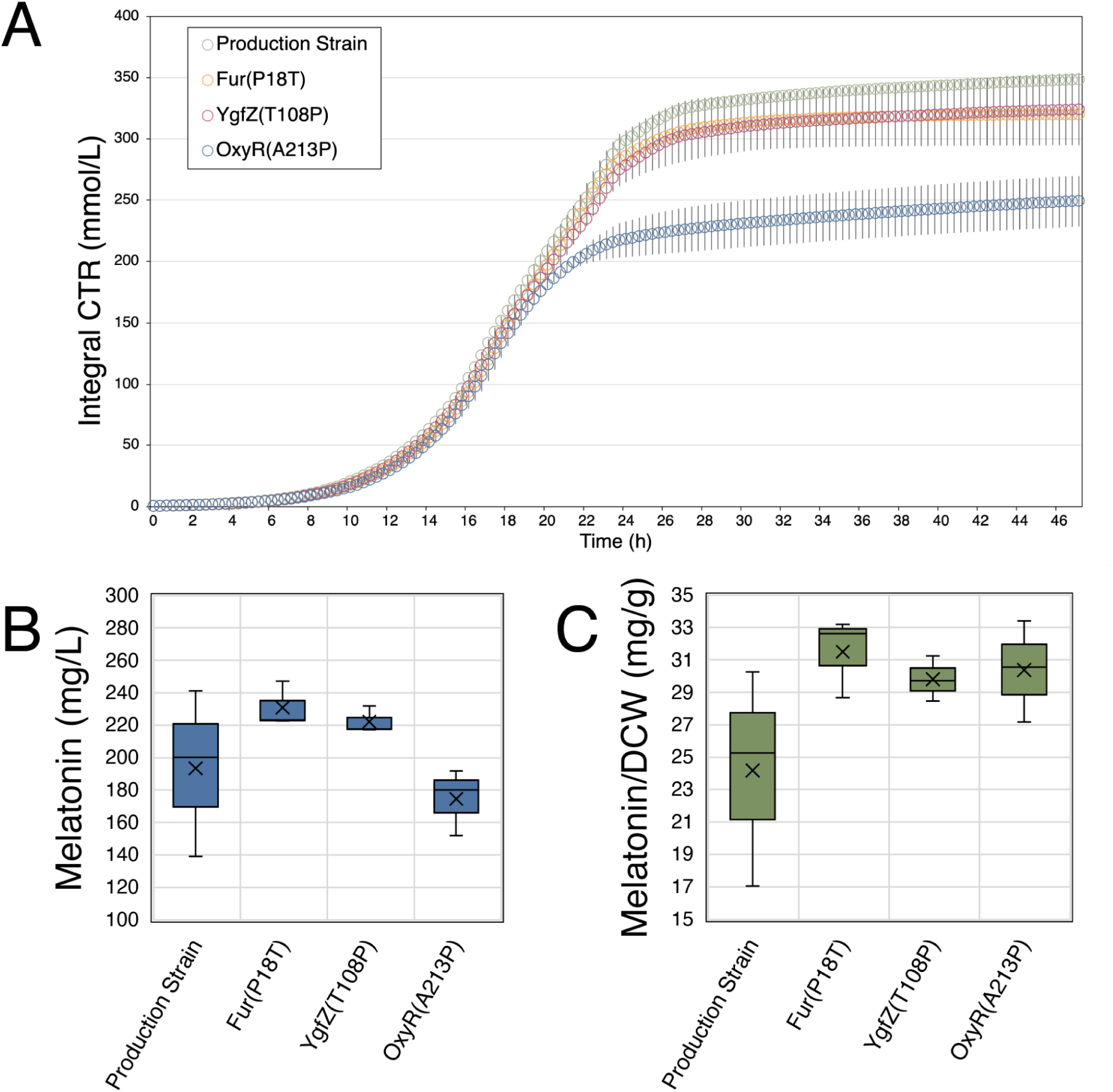
Batch fermentation of selected strains. We performed batch cultivation of the parent production strain (control), and strains with one of the 3 mutations incorporated: Fur(P18T), YgfZ(T108P) or OxyR(A213P). A) Growth curves represented by integral carbon dioxide transfer rate (CTR) measured online. Data represents the average of 3 replicates. Error bars indicate standard deviation. B) Melatonin final titers measured by HPLC after 48 h. C) Specific melatonin production of 4 strains. Dry cell weight (DCW) was converted from final OD (600 nm) measured manually by multiplying 0.341 (Abs/g_DCW_/L). Melatonin production is well-maintained in all the mutants. We did not observe statistically significant improvement (p>0.05) of the melatonin titer in mutant strains compared to the parent. The batch fermentation was performed using 6-deep well microplates (Enzyscreen, Heemstede, Netherlands), and incubated at 30°C with 225 rpm for 24 h using Kuhner TOM shaker (Kuhner, Birsfelden, Switzerland). The medium was modified from (27) and pH was adjusted to 6.5. HPLC was performed as described previously (48).

## Abbreviations

ALE: Adaptive Laboratory Evolution
ROS: Reactive Oxygen Species
WT: Wild-type
AA: amino acid

## Author Information

### Corresponding Author

Patrick Phaneuf - Novo Nordisk Foundation Center for Biosustainability, Technical University of Denmark, 2800 Kgs. Lyngby, Denmark

Lei Yang - Novo Nordisk Foundation Center for Biosustainability, Technical University of Denmark, 2800 Kgs. Lyngby, Denmark

### Author Contributions

Conceptualization and computational analysis: LY, PVP, and KR. Computational data processing: FB. Conceptual support: BOP. Designed selection experiment, troubleshot experimental data: LY, SHK. Strain engineering: SHK and CR. Executed selection experiment, and library preparation for sequencing:SHK. Writing: PVP, LY, SHK, and KR.

## Acknowledgements

The authors gratefully acknowledge Christina Lenhard and Suresh Sudarsan for their technical support and Emre Ozdemir for comments on the manuscript. We thank Mariana Arango Saavedra, Line Sondt-Marcussen, Arsenios Vlassis, and Vijayalakshmi Kandasamy from CfB-Biofoundry for helping with genome sequencing.

